# A Transfer entropy-based methodology to analyze information flow under eyes-open and eyes-closed conditions with a clinical perspective

**DOI:** 10.1101/2022.12.20.521265

**Authors:** Juan F. Restrepo, Diego M. Mateos, Juan M. Díaz López

**Affiliations:** Instituto de Investigación y Desarrollo en Bioingeniería y Bioinformática, CONICET–UNER, Entre Ríos, Argentina; Consejo Nacional de Investigaciones Científicas y Técnicas (CONICET), Argentina; Facultad de Ciencia y Tecnología. Universidad Autónoma de Entre Ríos (UADER). Oro Verde, Entre Ríos, Argentina; Instituto de Matemática Aplicada del Litoral (IMAL-CONICET-UNL), CCT CONICET, Santa Fé, Argentina; Facultad de Ciencias Químicas (FCQ), Universidad Nacional de Córdoba (UNC), Córdoba. Argentina; Instituto Argentino de Ciencias de la Conducta (IACCo). Córdoba. Argentina; Facultad de Matemática, Física, Astronomía y Computación, (FAMAF), UNC. Córdoba. Argentina

**Keywords:** Information Flow, Transfer Entropy, Eyes-open and Eyes-closed Resting States, Brain Dynamics, EEG

## Abstract

Studying brain dynamics under normal or pathological conditions has proven to be a challenging task, as there is no unified consensus on the best approach. In this article, we present a methodology based on Transfer Entropy to study the information flow between different brain hemispheres in healthy subjects during eyes-open (EO) and eyes-closed (EC) resting states. We used an experimental setup that mimics the technical conditions found in clinical settings and collected data sets from short records of 24 channels electroencephalogram (EEG) at a sampling rate of 65 Hz. Our methodology accounts for interhemispheric and intrahemispheric information flow analysis in both conditions and relies on 4 indexes calculated from the transfer entropy estimations between EEG channels. These indexes provide information on the number, strength, and directionality of active connections. Our results suggest an increase in information transfer in the EC condition for the alpha, beta1, and beta2 frequency bands, but no preferred direction of interhemispheric information movement under either condition. These results are consistent with previously reported studies conducted with denser EEG recordings sampled at a higher rate. In conclusion, our methodology shows a significant difference in the brain’s dynamics of information transfer between EO and EC resting states, which can also be applied to regular clinical sessions.

## 1 Introduction

The activity recorded by electroencephalography (EEG) usually comes from the encephalic cortex (brain and cerebrum). Once the inputs from the thalamus reach the fourth layer of the cerebral cortex, local microcircuits are responsible for amplifying and computing these signals. However, 75% of the cerebral cortex is composed of association areas that receive multiple multimodal inputs even under conditions of wakefulness, eyes closed or open [1]. This generates an activity known in clinical practice as basal activity. These dynamics are studied for diagnostic purposes. The strength of EEG lies in the electrophysiological dynamics concerning time in the order of seconds-minutes-hours, unlike evoked potentials (order of milliseconds). One of its limitations is its low spatial resolution (in the order of centimeters) compared to the precision achieved with functional Magnetic Resonance Imaging (fMRI). Even so, it is possible to describe local behavior in the order of centimeters -this can be seen in focal dysfunctions in epilepsy, mostly those with clinically observable partial seizures.

For the study of EEG dynamics, it is important to understand the collective behavior of the signals. In other words, the interrelationship that exists between the different channels. From this arises the concept of “functional connectivity” (FC), which can be defined as the statistical relationship between specific physiological signals over time. Beyond the existence of FC between two brain areas, there is no direct structural correlation observable by tractography that can demonstrate that these two brain areas are connected. It is possible to think that this FC would describe the shared dynamics between areas (similar functioning or shared information). This is the basis of the amplification and computation of neuronal signals, in the tangential component of axons connecting the cerebral cortex of the association areas—layers 1, 2, and 3. This micro connectivity cannot be studied by brain tractography so far, as it describes predominantly the behavior of projection fibers. For this reason, the study of FC is one of the most important areas to be studied in neuroscience. Previous studies showed a reduction in the FC during the eyes-closed condition compared to the eyes-open condition, this phenomenon is termed “alpha desynchronization” [2, 3]. Geller and colleagues found that closing the eyes causes generalized low-frequency power increase and focal gamma attenuation in the human electrocorticogram [4]. Barry et al. found topographical and spectral-power differences, in EEG recordings, between the eyes-closed and eyes-open resting states. Other work using the lagged linear coherence algorithm, as a measure of functional connectivity, found that the difference between closed and open eyes can be observed mainly in the alpha band, being larger in closed eyes, while the opposite is true for the beta1 and beta2 bands, which are larger in open eyes. A recent study, conducted on intracranial EEG data, found that phase-based connectivity is sensitive to the transition from eyes-closed to eyes-open condition, only in interhemispheric and frontal electrodes [5]. Other studies in EEG using symbolic analysis showed that this technique can distinguish between the two conditions [6]. In a statistical mechanic approach over the connectivity study, Guevara et al. gave evidence that in the baseline state, entropy increases during the eyes-open condition compared to the eyes-closed condition [7].

Nevertheless, the possibility of knowing how much information two EEG signals share has always been a challenge. Currently, there are several metrics to study the correlation between EEG signals, such as spectral coherence [8, 9], magnitude squared coherence [10, 11], amplitude envelope correlation [12, 13], synchronization probability [14, 15], phase lag index [16, 17], symbolic weighted mutual information [18, 19], however, these measures do not give information about the direction of the information flow, i.e. which channel is the source and which one is the target. To address this problem, other metrics that measure the directionality of information have been developed, e.g., Granger causality [20] or transfer entropy (TE) [21].

Transfer entropy is a measure of information transport based on Shannon entropy [21]. This measure can quantify the strength and direction of the coupling between simultaneously observed systems [22]. Moreover, it has attracted general interest because it can be used to study complex interaction phenomena found in several disciplines [22]. In particular, we can mention some of the applications of TE in different areas of neuroscience. For example, Olejarczyk et al. used transfer entropy, multivariate TE (also called partial TE), and other measures to analyze effective connectivity in resting state EEG data. They recruited 19 healthy subjects and recorded a 128-electrodes EEG with a sampling frequency of 1 kHz and using the 10-10 international system, and the results suggest that the biggest difference in TE between eyes conditions (closed and opened) was given in the alpha band [23].

Bagherzadeh et. al. presented a method for schizophrenia detection, which involves building a connectivity matrix based on TE and feeding it to a deep-neural network (different deep-learning models were investigated). The study included 14 healthy individuals and 14 schizophrenia patients, from whom a 19-channels EEG was collected under the eyes-closed condition at a sampling rate of 250 Hz (using the 10-20 system). The reported accuracy and F1-score were both above 99.9%, and the study’s findings indicate that normal individuals exhibit higher transfer entropy values (on average) compared to schizophrenia patients. Finally, the authors discuss two limitations of their approach: a small sample size for training the network and the high computational cost of fine-tuning [24].

In [25] the authors combine a dyadic stationary wavelet decomposition with the TE to assess cortico-muscular information flow. The idea was to analyze the coupling between EEG and electromyogram signals within a time-scale decomposition approach. Their results suggest that EEG to EMG information flow is observed mostly in alpha and beta bands but its strength and directionality depend on the observed scale.

Another study analyzes the changes in the brain neurodynamics among resting-state before and after working memory processes (encoding and retrieval phases) [26]. They obtained the EEG (60 channels) from 29 participants at a sampling rate of 1 kHz and analyze changes in power and information flow (through phase TE) between resting state periods. Their conclusions state that there are differences between these resting states in the delta, alpha, and beta bands. Moreover, they suggest that power and information flow in the alpha band during the resting state are important markers related to working memory process performance.

Zhang et. al. used multivariate TE over magnetoencephalographic signals to characterize subjects suffering from depression and healthy controls. They suggest that this information flow measure was generally lower for the depression group compared to the healthy group, moreover, the biggest difference between populations can be observed in the gamma band (*>* 30 Hz) of the frontal region [27].

Finally, a more recent article suggests that phase TE of the delta, theta, alpha, and beta bands can be used as biological markers to distinguish between subjects with prolonged disorder of consciousness and healthy ones [28].

All these studies have something in common, their protocols include high-density EEG recorded at high sampling rates. These conditions are difficult to meet in a clinical environment, which is characterized by using short-time and low-density EEG recordings, with a low sampling rate.

In this work, we analyze 15 EEG recordings from different subjects during eyes-closed and eyes-open resting states following a TE framework but with a clinical perspective. The characterization of the information flow between cortical areas in resting conditions is fundamental for describing patterns observed in control subjects. This would give us a baseline to study connectivity changes in pathological scenarios.

The contribution of this article can be listed as follow:

- We have proposed a method to analyze information flow between EEG channels during the open eyes and closed-eyes conditions in healthy subjects.
- The methodology has been designed taking into account the times and technological limitations presented in a general clinical consult.
- Our results suggest an increased information transfer in the closed eyes condition for the alpha, beta1, and beta2 frequency bands.

The structure of the article is the following: Section II outlines the participant recruitment criteria, data acquisition, and preprocessing, as well as presents the proposed methodology to analyze the information flow. Section III discusses the study’s results, which include a comparison of information flow in different frequency bands between eyes-open and eyes-closed conditions. In Section IV, our results are compared with those obtained by similar studies, followed by a discussion of their clinical implications. Finally, Section V summarizes the study’s key findings, as well as suggests further research using this methodology.

## 2 Methods

### 2.1 Participant recruitment

This study has a comparative, observational, prospective design between two electrophysiological conditions (open and closed eyes) in the same control group during the resting state. The 15 subjects were selected using the following inclusion criteria: a) Women and men aged between 20 and 50 years old. b) Individuals holding 7 or more years of complete formal education. c) Individuals without neurological and psychiatric medical history and normal values on the Beck depression inventory [29] and State-Trait anxiety inventory, form X and Y [30]. d) Individuals with a normal EEG report according to St. Louis EK and Frey LC’s [31] criteria of normality. e) Not to consume psychotropic drugs or drugs with effects on the central nervous system. f) Not to present an abusive consumption of psychoactive substances such as cocaine, marijuana, alcohol, or others. g) Not to have had a moderate or minimum consumption of alcohol in the 48 hours prior to the full investigation evaluation. h) Individuals that have accepted and signed the informed consent. This research protocol was accepted and approved by the ethics committee of the Hospital Córdoba (Córdoba-Argentina) and registered in the Ministry of Health of the Córdoba Province on 20-03-2017 under number 3170.

### 2.2 EEG recordings and preprocessing

The EEG data were measured according to the standards of the International Federation of Clinical Neurophysiology [32], following the international 10-20 system. The recordings have monopolar montages, with a bimastoid reference. Twenty channels (F1-F3-C3-P3-O1-F7-T3-T5-Fz-Cz-Pz-Oz-T6-T4-F8-O2-P4-C4-F4-F2) were recorded with a 24 channels polysomnograph (Neutronic, model ME-2400) at 65 Hz sampling frequency.

EEG channels were average referenced and filtered in the following frequency bands: [4, 8) Hz (theta), [8, 12) Hz (alpha), [13, 19) Hz (beta1), [19, 25) Hz (beta2) and [25, 31] Hz (beta3) [33]. The instantaneous phase of each EEG channel was estimated through the Hilbert transform [23].

### 2.3 Transfer entropy estimation

Transfer entropy [21] between all pairs of EEG electrode phases, was calculated (in both directions) with the k-nearest-neighbor (KNN) methodology proposed in [34]^1^. Phase signals were embedded using an embedding-dimension equal to 4 and an embedding-lag that changes in accordance with the frequency band (*τ* = 10 for theta band, *τ* = 8 for the alpha band, *τ* = 7 for beta1 band; and *τ* = 6 for beta2 and beta3 bands). Each value of *τ* was obtained as the first zero of the correlation function between the filtered phases of the F1 and F3 channels (very close and correlated channels). For the KNN algorithm, the number of the nearest neighbors was set to 8.

A conventional surrogate test was chosen to evaluate the statistical significance of the TE estimations [35]. Here, the null hypothesis is that there is no relation between the target and the source signals, i.e. *Txy* = *Tyx* = 0. Two different surrogate distributions were calculated for each TE estimator. The first surrogates were estimated from circular-shifted versions of the original phases. A random index was sampled from a uniform distribution, and both signals (target and source) were shifted with the same index. These surrogates preserve all the characteristics of the original phases, as well as the time connection between the target and the source. However, the estimation of TE differs from the original data, because of the embedding scheme used by the KNN algorithm. The estimations from these surrogates generate a distribution grouped around the value calculated from the original phases and account for the variability of the algorithm. The second surrogates seek to break any possible link between source and target signals while keeping each one inner correlation. A surrogate realization of the source was obtained using the inverse Fourier transform of the product between the amplitude spectrum of the signal and the Fourier phase of random normal noise [36]. Then, a surrogate estimation of TE is calculated from the target signal and the surrogate source.

To analyze the statistical significance of the TE estimation between a pair of channels’ phases, we used the following methodology: we compute the difference *T* =*< Txy − Tyx >* where the median value is estimated along the first surrogate realizations (including the one with the original data). Then, we obtain the distribution of the difference between Fourier surrogates 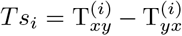, where 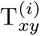 is the i-th Fourier surrogate of *Txy*. Finally, we perform a two-sided statistically significant test, using the difference *Tsi* as surrogate distribution and *T* as test statistics. The significance level was set to *α* = 0.05, and we computed 512 realizations of each surrogate. If the probability of the test statistic is greater or equal to 1 − *α/*2, we consider that the difference *T − T* is statistically greater than zero and *T > T*. On the other hand, if the probability of *T* is below or equal to *α/*2, we conclude that *T*_*xy*_ < *T*_*yx*_. Otherwise, we consider that *Txy − Tyx* = 0.

Following the above-mentioned procedure (see figure 2), we construct two matrices named the transfer entropy matrix (Tent matrix) and the flow matrix (figure 1). For a single subject and a given frequency band, the transfer entropy matrix (figure 1a) shows the difference *T*_*xy*_*−T* _*yx*_ between EEG channels. A positive value (*T*_*xy*_ *> T* _*yx*_) indicates that information is greater in the *X→Y* direction than in the opposite. For example, the red-shaded value in the 2nd-row and 7th column says that the information flows from *F* 3 channel to the *T* 3 channel. The flow matrix (figure 1b) can be read in the same way, however, it contains only three values [−1, 0, 1] given by sgn(*T*_*xy*_*−T*_*yx*_). In this sense, the flow matrix gives us information about which connections, between EEG channels, are active; whereas the transfer entropy matrix tells us the amount of information shared through that connection. Note these matrices are upper triangular since *T*_*xy*_*−T*_*yx*_ = − (*T* _*yx*_ *−T* _*xy*_). Finally, we divided the area of these matrices into smaller subareas (figure 1c), each of them representing a group of connections between EEG channels. For example, the matrix subarea *G*3 contains the connections between the channels placed over the left hemisphere and the channels placed over the right hemisphere.

**Figure 1.**
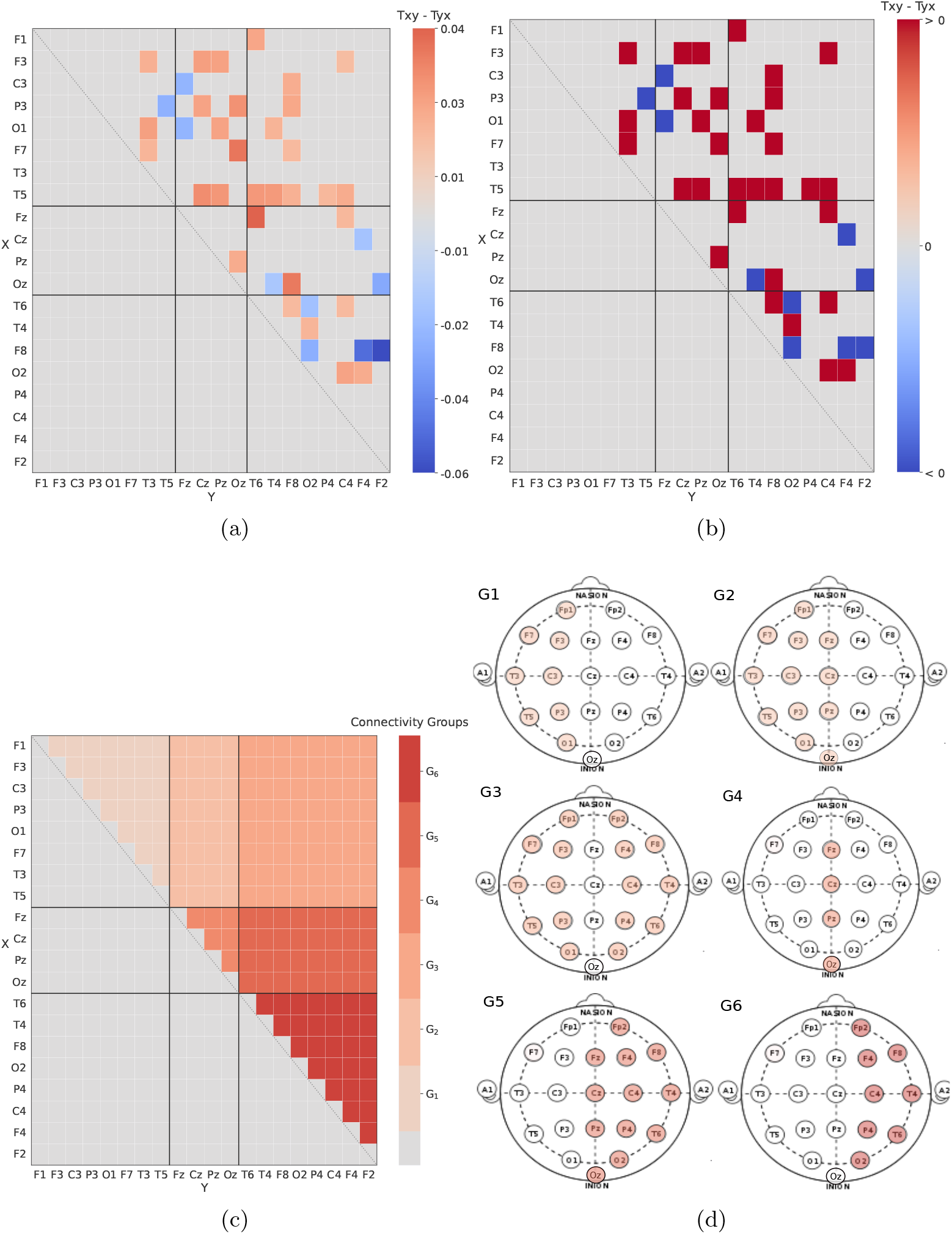
Information flow matrices of a single subject in the alpha band: (a) Transfer entropy matrix (Tent matrix). (b) Flow matrix. (c-d) Connectivity groups.

**Figure 2.**
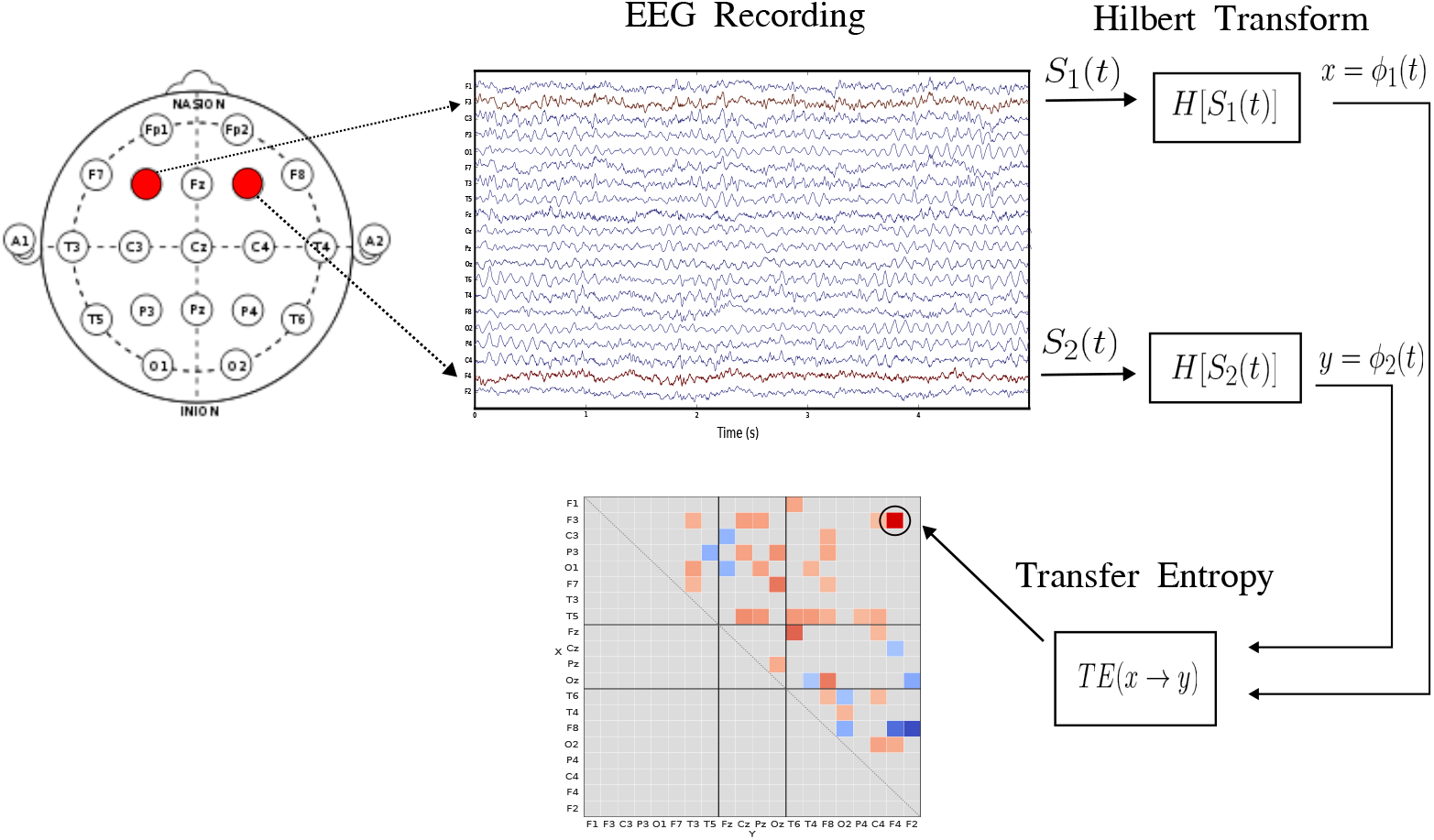
Transfer entropy matrix estimation methodology.

From the information provided by these matrices we define a set of measures over each subject (table 1): The Total Activity is calculated, from the information flow matrix, as the sum of all active connections between channels over all possible pairwise (190), it is an estimation of the number of interactive channels. The Total Tent (Total Transfer Entropy) is the sum of transfer entropy values over the active connections, and it accounts for the magnitude of the information transferred between active channels. The Local Activity measures the number of interactive channels in each connectivity group and is computed as the number of active channels over all possible connections inside that group. Please note, each group, shown in figure 1c, has a different number of possible connections. Finally, the Directed Activity measures the proportion of activity that goes in a particular direction (*X* →*Y* or *Y* → *X*), which is calculated as the sum of active connections in that direction over the total number of active connections.

**Table 1:**
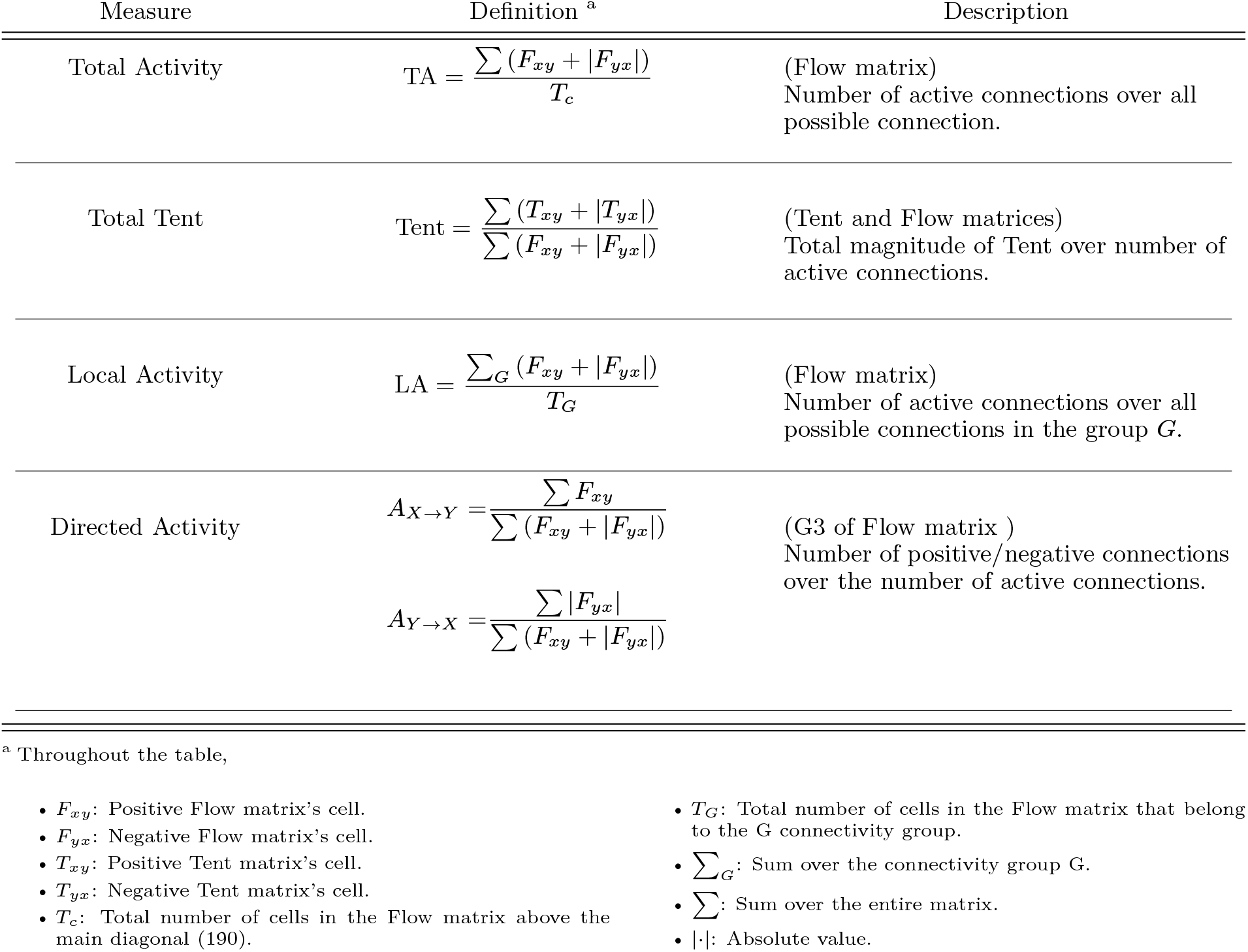
Measures definition over the Flow and the Transfer Entropy matrices

## 3 Results

In order to analyze the Total Activity between different eyes conditions, we have applied a Wilcoxon signed-rank test for each frequency band (figure 3a). Moreover, we have used a Conover-Iman test with a Holm-Bonferroni correction and a *H*0 rejection threshold of 0.05 to verify if there are differences for each eyes’ condition between the frequency bands (see table 2).

**Table 2:**
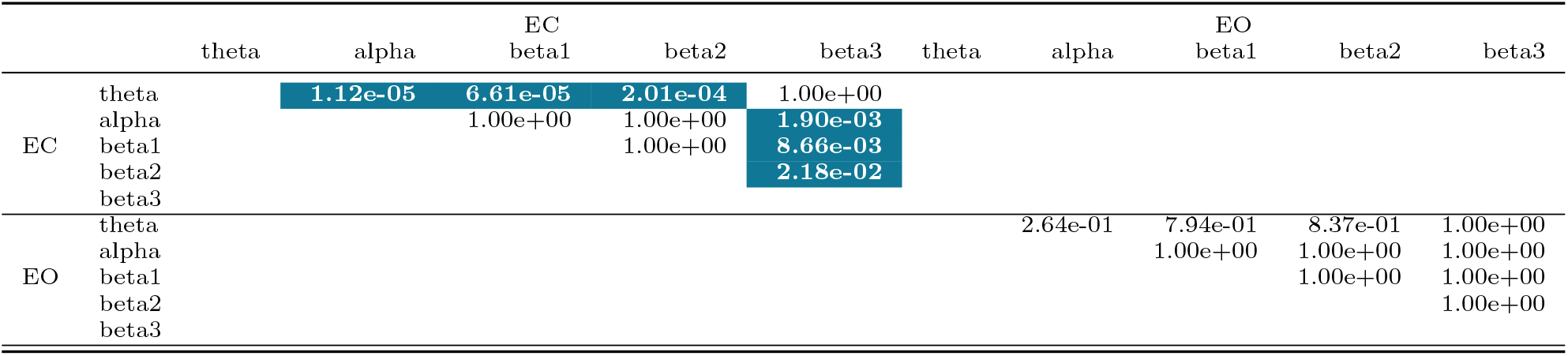
Total Activity Conover–Iman test p-value for different frequency bands and same eyes condition.

**Figure 3.**
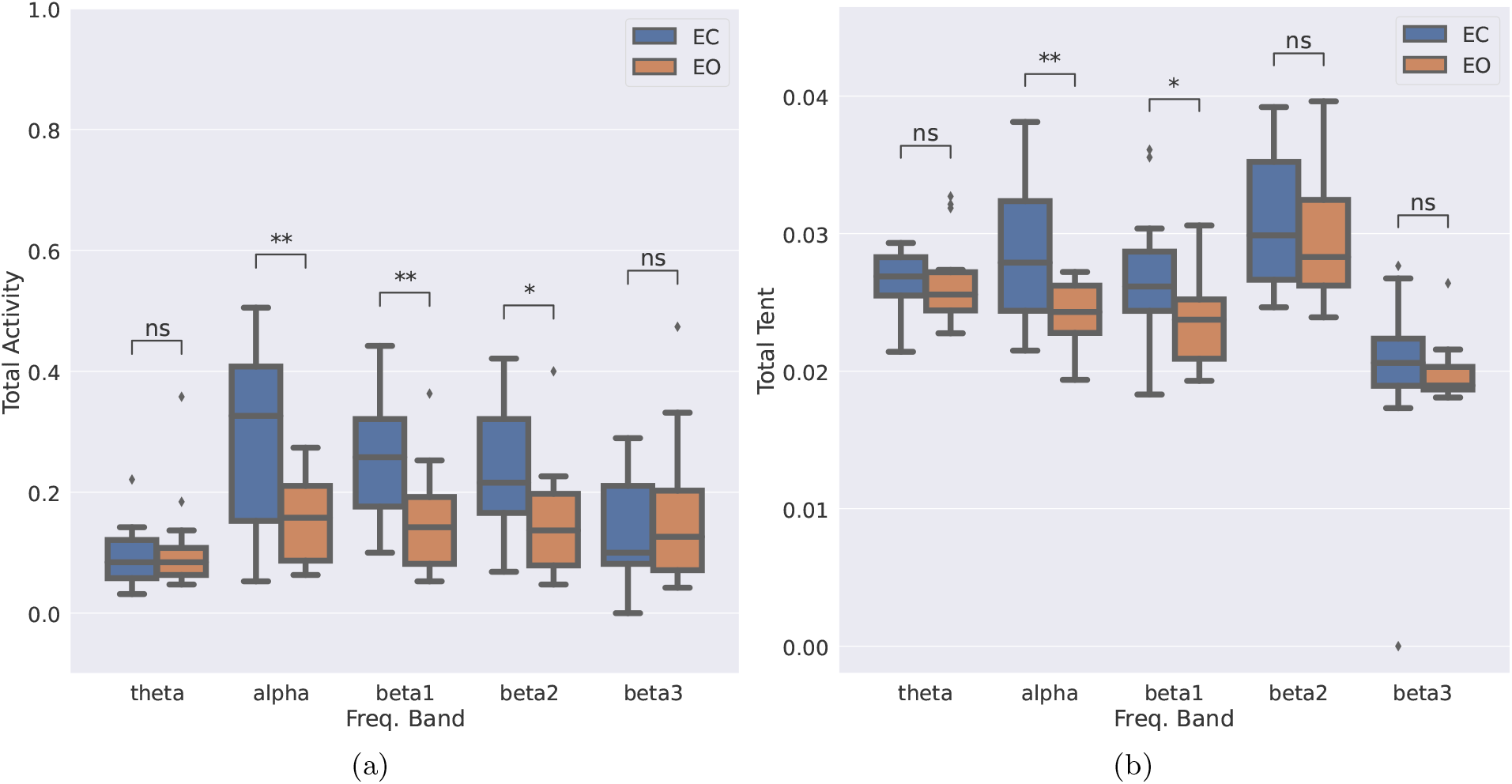
(a) Total Activity. (b) Total Tent. Wilcoxon signed-rank test p-values range: ns: *p ≤* 1, *\**: 0.01 *< p ≤* 0.05, *\*\**: 0.001 *< p ≤* 0.01.

The magnitude of the information transferred for different frequency bands and eyes conditions can be evaluated through the Total Tent estimator. As well as in the above analysis, we have made a Wilcoxon signed-rank test for each frequency band to evaluate differences between the two eyes conditions (figure 3a), and a Conover-Iman test for differences across the frequency bands (see table 3).

**Table 3:**
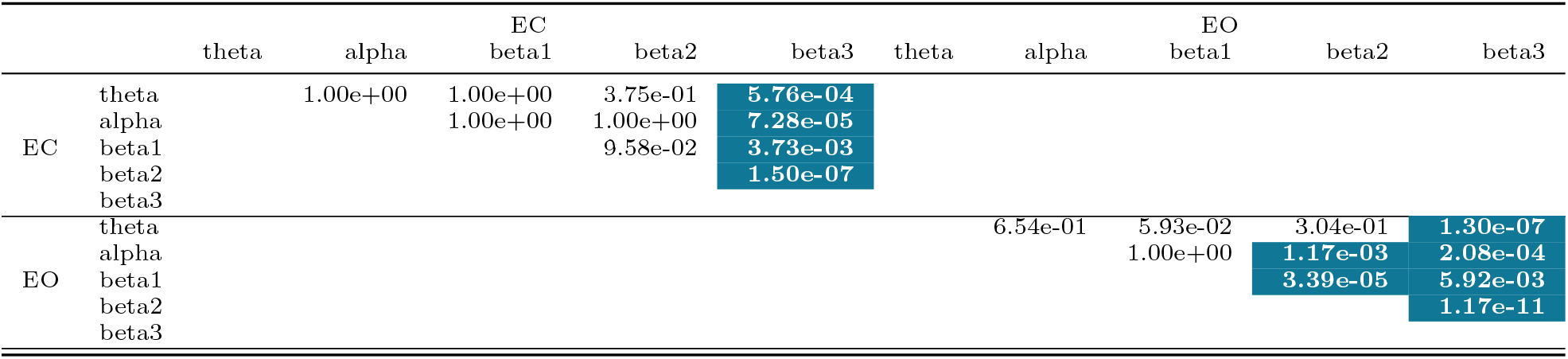
Total Tent Conover–Iman test *p*-value for different frequency bands and same eyes condition

Figure 3a shows the Total Activity values of the EO and EC conditions for the different electrophysiological bands analyzed. The Wilcoxon test shows greater activity for the EO condition in alpha, beta1, and beta2 bands. Moreover, figure 3b presents the Total Tent values for the different conditions and bands. As can be seen, only alpha and beta1 bands show significantly higher values for EC than for the EO condition.

We performed a Conover–Iman test to analyze if there are differences in Total Activity (table 2) and Total Tent (table 3) across frequency bands for each condition. From table 2 we do not observe any difference in the number of active connections for the EO condition, but for the EC condition, there is less activity in the theta and beta3 bands. Table 3 shows that the beta3 band has less amount of transferred information in both eyes conditions.

The Local Activity for different connectivity regions^2^ in the analyzed frequency bands can be seen in figure 4. For the theta and beta3 bands (figures 4a and 4e) we do not detect any difference between eyes conditions. On the other hand, we observe a greater number of active connections in the EC condition for the alpha, beta1, and beta2 bands in almost all connectivity groups (see Figures 4b, 4c and 4d). This result is under the Total Activity values.

**Figure 4.**
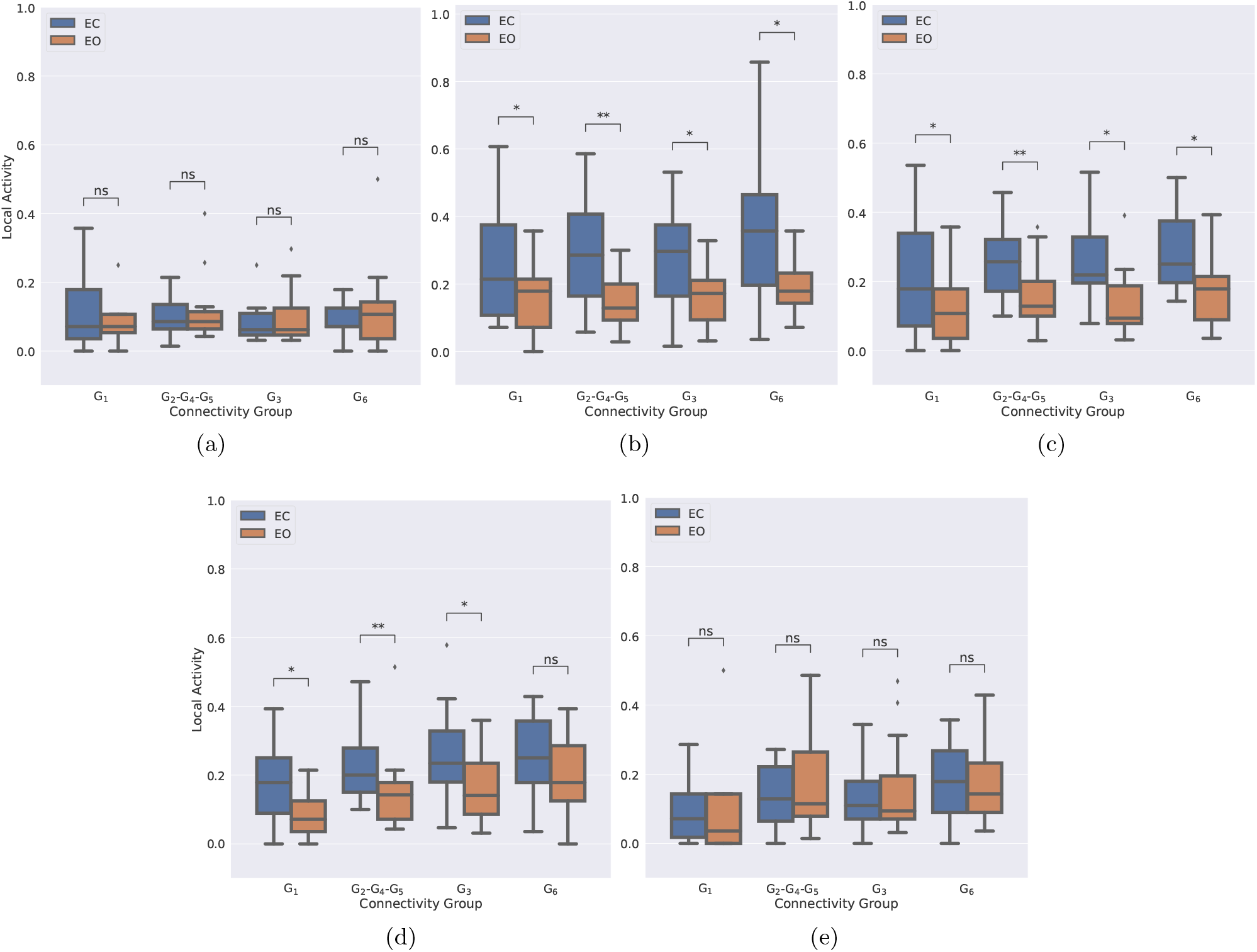
Activity by regions: (a) theta band. (b) alpha band. (c) beta1 band. (d) beta2 band. (e) beta3 band. Wilcoxon signed-rank test *p*-values range: ns: *p ≤* 1, *\**: 0.01 *< p ≤* 0.05, *\*\**: 0.001 *< p ≤* 0.01.

Since the *G*3 region represents the interhemispheric information flow, we studied the direction of the flow in the different conditions and bands. Figure 5 shows the Directed Activity for EC and EO. For both conditions, none of the bands present significant values, suggesting that there may not be a preferred direction but a bi-directionality of the information flow.

**Figure 5.**
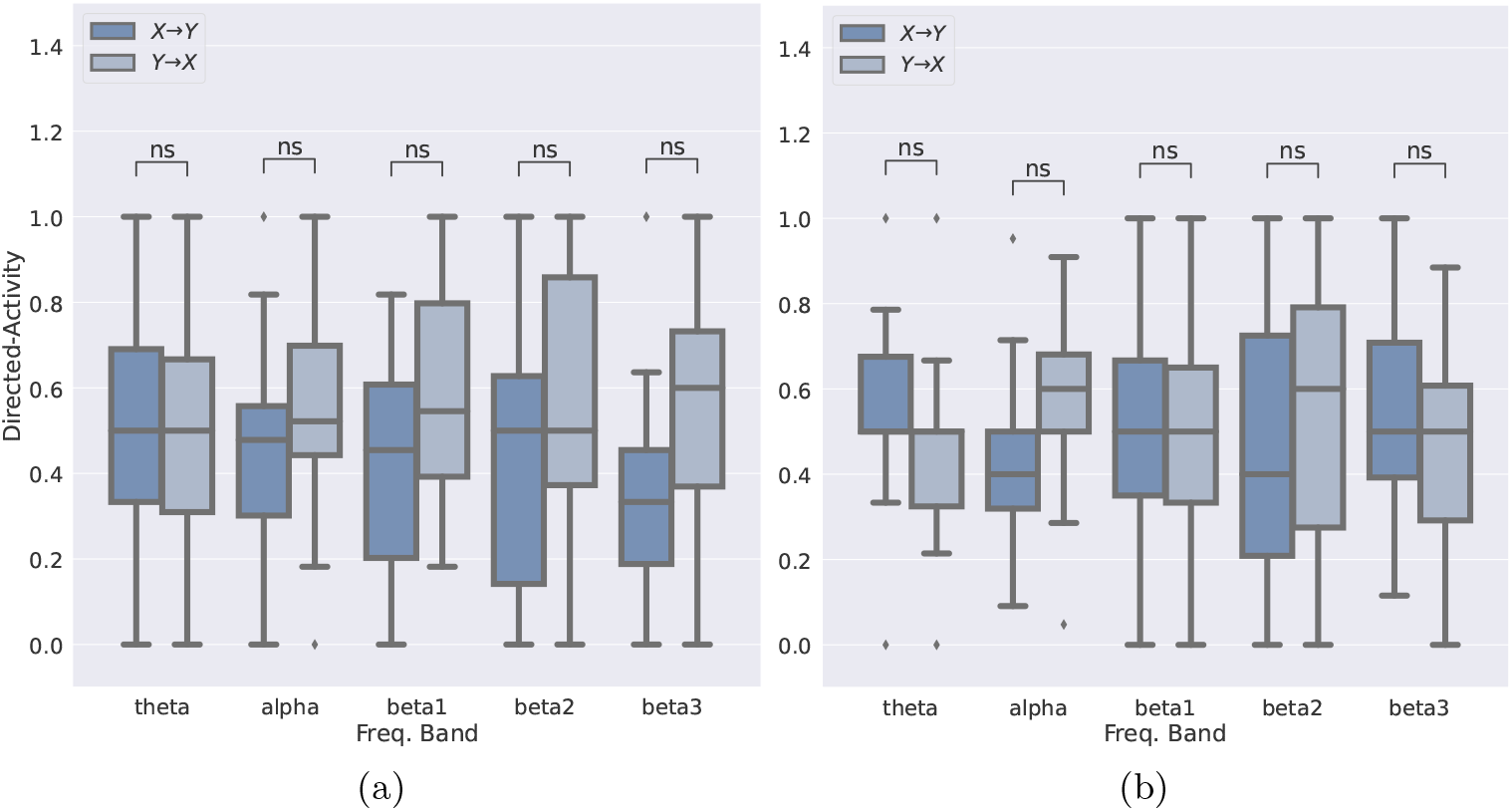
Connectivity group G3 Directed Activity: (a) Flow direction EC. (b) Flow direction EO. Here *X* and *Y* represents the left and right hemispheres, respectively. Wilcoxon signed-rank test *p*-values range: ns: *p ≤* 1, *\**: 0.01 *< p ≤* 0.05, *\*\**: 0.001 *< p ≤* 0.01

## 4 Discussion

The purpose of this study is to characterize the information flow during EO and EC conditions in healthy subjects. In this sense, we seek to evaluate some of its aspects, such as the number of active connections, the magnitude of the information transferred through those connections, and the flow direction. Moreover, the difference in these quantities by frequency bands and the contribution of each connectivity group to the global information flow.

The idea of “functional connectivity” raises some controversy because of the relationship between its conceptual imprecision and the multiplicity of meanings assigned to it in the literature, as a result of the observation of multiple physical phenomena. Sometimes it is used to account for temporal changes in brain perfusion observed by fMRI, sometimes it is related to the measure of coherence or correlation between EEG and/or MEG signals describing voltage variation over time. Recent studies attempt to establish some kind of correlation between the two approaches, referring to Functional Neuroimaging studies, those that describe the functional organization of the brain independently of the observation instrument [37].

Even if one studies a phenomenon through a single observation instrument, e.g., EEG, this does not guarantee that there is a single definition of functional connectivity. In our case, the idea of functional connectivity is given by the information that is shared or transferred between 2 EEG signals, defined by the bivariate TE. From this measure, we calculate two matrices, one account for the active connections (flow matrix), and the other accounts for the magnitude of the information flowing by those connections (transfer entropy matrix); then over these matrices, we calculate four information quantifiers (Total Activity, Total Tent, Local Activity and Directed Activity) which allow us to extract different and complementary information from each one. It is worth mentioning that, due to the lack of independence between observations (EEG records) in our dataset and the information quantifiers not following a known distribution, the usage of the ANOVA statistical test was not feasible. As a result, we chose to utilize the Wilcoxon signed-rank test to determine differences between eyes conditions and the Conover-Iman test to evaluate differences between frequency bands.

Our findings reveal that during the EC condition, the Total Activity (number of active connections) is greater than during the EO condition in the alpha, beta1, and beta2 bands. The most significant differences are observed in the alpha and beta1 bands, which is consistent with the results presented in [23] where the authors reported a higher disparity in TE between eye conditions in the alpha band. Their study consisted of the EEG analysis of 19 healthy young subjects, acquired through 128 electrodes at a sampling rate of 1000 Hz. They calculated an adjacency matrix (which is the same as our transfer entropy matrix) and obtained indexes based on graph theory. It is noteworthy that we obtained similar results through a simplified and more straightforward protocol setup (fewer EEG channels and reduced sampling frequency) and the indexes proposed by us are easier to interpret and calculate.

In a recent study by Song et al. [38], the researchers examined information flow changes during the transition between sleep stages. To do this, 40 healthy participants were enlisted and underwent a polysomnography examination, recording a 16-channel EEG using the 10-20 system at a 500 Hz sampling rate. The phase TE was calculated between pairs of EEG channel phases derived from a Morlet wavelet, as opposed to the Hilbert transform employed in this article. Additionally, the authors introduced two anisotropy indexes to describe the information flow across brain regions. Among various interesting findings, they observed significant differences in delta and alpha bands when transitioning from wakefulness to sleep stage N1. They also suggest a dynamic balance between the hemispheres, which aligns with our results as we did not observe a preferred direction for interhemispheric information flow.

Ekhlasi et al. assessed information flow direction between brain regions in children with attention deficit hyperactivity disorder (ADHD) and healthy participants [39]. A total of 121 children participated, with 60 being healthy individuals. The EEG aggregate data were collected using 19 channels and a sampling rate of 128 Hz. Their phases were obtained using the Hilbert transform, and the TE was calculated using histograms which we decided to replace with a KNN method. The authors proposed 4 indexes to characterize information flow accounting for the transferred info within the anterior and posterior regions, between them, and between the left and right hemispheres. In this study, we did not examine the information flow within the anterior and posterior regions. Nevertheless, our indexes can be easily adjusted to facilitate the examination. Although they measured the flow of information between both hemispheres in both directions, they didn’t indicate whether the flow of information was asymmetrically biased toward one direction. However, they observed that there was no significant difference between the groups with ADHD and healthy participants in terms of inter-hemisphere interaction at any frequency band.

From table 2 we see there is no statistical difference between frequency bands alpha and beta1 within the same physiological condition. This may suggest that the number of active connections could be independent of these frequency bands. The questions about why these frequency bands are predominant? Why there are more active connections in EC condition? and what are the intrinsic neurophysiological mechanisms that produce this response? Can not be answered by the present experimental setup.

Using transfer entropy versus the use of multivariate transfer entropy to study brain dynamics is a topic of wide discussion. In [23] the authors compare both information transfer measures, concluding the transfer entropy is susceptible to false positive connections. The multivariate TE is a measure employed to avoid indirect information transfer. For example, suppose, there are 3 random processes with the following causation chain: *X→Y→Z*; here, the TE will confirm that *Y* causes *Z* but will also say that *X* causes *Z*, which is a false link since *X* is not connected directly with *Z*. The multivariate TE was designed to avoid these kinds of false positive connections. However, increasing the number of variables entails a bigger increase in the data needed for its valid calculation. This condition is very difficult to satisfy in a clinical context where the time of signal acquisition is limited. In our experimental setup, we choose to calculate the transfer entropy in conditions that are met in the clinic, such as 20 to 32 EEG channels setup, 65 Hz as sampling frequency, and 30-second signal length.

Despite some limitations, such as a limited number of patients and a low sampling rate of the equipment, it is worth noting that the sampling rate in daily clinical practice is typically lower than in research settings. Nevertheless, our study yielded results similar to those found in studies with more complex equipment setups and a bigger number of subjects.

## 5 Conclusions

We have proposed a simple approach to analyze information flow between EEG channels during open and closed eyes conditions in healthy subjects. Our approach is based on TE quantifiers that can be computed on general clinical consults. Our findings reveal a greater number of active connections and increased magnitude of information transfer during the closed eyes condition, in the alpha, beta1, and beta2 frequency bands, compared to the open eyes condition. This transfer entropy approach can be useful for studying different electrophysiological conditions in clinical practice. Our aim for future works is to demonstrate the robustness of the methodology for different datasets and ensure the consistency of information transfer in the same electrophysiological condition by using experimental designs with repeated measures. In addition, we plan to investigate entropy transfer in hyperventilation and opto-stimulation conditions and analyze information flow in both resting and activity conditions through a standardized neurocognitive assessment method.

## Author contribution

**Juan F. Restrepo:** Conceptualization, Methodology, Software, Data curation, Visualization, Formal Analysis; **Diego M. Mateos:** Software, Visualization, Formal Analysis, Validation; **Juan M. Díaz López:** Conceptualization, Methodology, Resources. All authors participated in writing and editing this article.

## Aknowledgements

This work was partially by the Grant PICT 2019-01750 (ANPCyT), PICT 2020 SERIEA-01865 (ANPCyT), PIP 633 (CONICET) and PID 6228 (UNER).

## Footnotes

1 The code for Python or Matlab is located at https://bitbucket.org/jrinckoar/knn-informationmeasures/src/master/

2 Regions *G*2, *G*4, and *G*5 were grouped because they represent the activity in the middle-line EEG channels.

## Notes

### Competing Interest Statement

The authors have declared no competing interest.

### Summary of Updates

We have added new information, included additional bibliography, and rewritten some paragraphs to make it more understandable.

